# Going with the flow – how a stream insect, *Pteronarcys californica,* exploits local flows to increase oxygen availability

**DOI:** 10.1101/2022.06.06.495063

**Authors:** Jackson H. Birrell, H. Arthur Woods

## Abstract

For insects, aquatic life is challenging because oxygen supply is typically low compared to air. Although many insects rely on stream flows to augment oxygen supply, oxygen limitation may occur when oxygen levels or flows are low or when warm temperatures stimulate metabolic demand for oxygen. Behavior may allow insects to mitigate oxygen shortages – by moving to cooler, more oxygenated, or faster flowing microhabitats. However, whether stream insects can make meaningful choices depends on: i) how much temperature, oxygen, and flow vary at microspatial scales in streams and ii) the ability of insects to exploit that variation. We measured microspatial variation in temperature, oxygen saturation, and flow velocity within riffles of two streams in Montana, USA. Additionally, we examined the preferences of nymphs of the stonefly *Pteronarcys californica* to gradients of temperature, oxygen, and flow in lab choice experiments. Temperature and oxygen level varied modestly within stream riffles (∼ 1.8 °C, ∼ 8.0% of air saturation, respectively). By contrast, flow velocity was highly heterogeneous, often varying by more than 125 cm s^-1^ within riffles and 44 cm s^-1^ around individual cobbles. Exploiting micro-variation in flow may thus be the most reliable option for altering rates of oxygen transport. In alignment with this prediction, *P. californica* nymphs showed relatively little ability to exploit laboratory gradients in temperature and oxygen. By contrast, they readily exploited micro-variation in flow – consistently choosing higher flows when conditions were warm or hypoxic. These behaviors may help stream insects mitigate low-oxygen stress from climate change and other anthropogenic disturbances.

**Summary Statement:** Stonefly nymphs stressed by higher temperatures and lower oxygen availability often relocate to areas of higher flow. This behavior likely increases the ratio of oxygen supply to demand.

## Introduction

Life in water is shaped by the scarcity of oxygen (Hutchinson, 1981; Lancaster & Downes, 2013). Water contains ∼ 30 times less oxygen than air (Jones et al., 1972). Water is also dense and viscous, which slows rates of diffusion and makes respiratory ventilation energetically expensive (Denny, 1993; Verberk & Atkinson, 2013). For water-breathing ectotherms like aquatic insects, the challenge is to extract sufficient oxygen to fuel metabolism, despite low levels of environmental oxygen supply (Verberk et al., 2016; Woods & Moran, 2020).

This oxygen problem is complicated further by variation in temperature and water velocity (Woods, 1999; Jacobsen, 2003; Verberk et al., 2011; Frakes et al., 2021; Verberk et al., 2016b). Warmer water modestly increases rates of oxygen supply but raise organismal demand even more (Woods, 1999; Woods & Moran, 2008; Verberk et al., 2011; Verberk & Atkinson, 2013). At high temperatures, oxygen demand can exceed organisms’ capacities to supply sufficient oxygen, resulting in declines in organismal performance and potentially death (Pörtner, 2007; Verberk et al., 2016; Harrison et al., 2018). Higher flows, by contrast, increase rates of oxygen supply by thinning boundary layers enveloping the insect and its respiratory surfaces (e.g., gills or thin segments of the cuticle) (Hynes, 1970). Water velocity is zero (i.e., no slip condition) at the cuticle-water interface but increases with distance from the body surface until it approaches the velocity of the free-stream flow (Vogel, 1981). When boundary layers are thin, as in fast-flowing and turbulent water, oxygen diffuses over shorter distances between the water column and respiratory tissues, increasing supply rates (Pinder & Feder, 1990; Hall & Ulseth, 2019). Thus, aquatic ectotherms can tolerate deeper hypoxia and warmer temperatures when water velocities are sufficiently high (Frakes et al., 2021).

Taken together, climate change and other anthropogenic disturbances, like eutrophication and dams, are raising water temperatures, depressing oxygen concentrations, and lowering water velocities (at least seasonally) in many rivers and streams. These changes threaten aquatic communities in part by depressing the ratio of oxygen supply:demand (Jacobsen et al., 2008; Deutsch, et al., 2015; Verberk et al., 2016b). Predicting how aquatic communities will respond, however, will require a better understanding of the mechanisms that aquatic insects use to mitigate oxygen limitation. A broad literature now addresses the mechanisms by which ectotherm thermal performances, thermal limits, and acclimation responses depend on oxygen and flow (Verberk et al., 2011; Verberk et al., 2016; Rubalcaba et al., 2020; Collins et al., 2021; Frakes et al., 2021). Much less literature has focused on behavioral mitigation of the oxygen problem (but see Verberk & Bilton, 2015), even though behavior is well known to provide small terrestrial and intertidal ectotherms with the means to thermoregulate (Ebersole & Frissell, 2001; Helmuth et al., 2007; Scheffers et al., 2014; Woods et al., 2015; Birrell et al., 2020; Pincebourde & Woods, 2020). Whether aquatic insects can mitigate oxygen shortages by moving among microclimates will depend on (i) whether diverse conditions are available locally and (ii) whether insects actually exploit that diversity, if available. We expect that these two conditions are related to one another – in the sense that aquatic insects should evolve behavioral mechanisms to exploit microclimatic mosaics only if they are exposed regularly to biologically meaningful gradients in nature. Assessing these points will be critical to understanding how aquatic insects will respond to future climate change and has so far received little attention (e.g., Helmuth et al., 2007; Pincebourde & Woods, 2020).

Which of the three factors we consider – temperature, oxygen, and flow – are likely to show strong local gradients? Because water has a high heat capacity, equilibrates slowly with local gas partial pressures, and is typically well-mixed (in lotic systems), we predict that microclimatic diversity in temperature and oxygen will be slight. Cool microclimates can form in association with groundwater inputs, side channels, springs, deep-water pools, and shading (Mosley, 1983; Matthews & Berg, 1996; Clark et al., 1999; Ebersole & Frissell, 2001; Ebersole et al., 2003), but the spatial grain at which these differences are manifested appears too coarse for individual insects to exploit (e.g., Ebersole & Frissell, 2001). Moreover, cold patches are often associated with inputs of hypoxic groundwater, which would at least partially counteract the benefits of such thermal refugia (Matthews & Berg, 1996; Torgersen et al., 1999; Elliot, 2000). Small gradients in temperature and oxygen may also form in stream riffles as a consequence of patterns of exchange between in-stream and hyporheic flows, with slightly warmer, more oxygenated water at riffle heads and slightly colder, less oxygenated water at the tail (from upwelling of hyporheic water) (Davy-Bowker et al., 2006).

By contrast, microclimatic diversity in flow velocities are often very high in lotic systems. In streams, flow velocities often vary from ∼ 0 within the substrate to > 100 cm s^-1^ in the free stream environment (White, 1990; Comiti et al., 2007), and individuals may often be able to alter local flows past their bodies by moving just a few cm. Indeed, gradients in flow velocity strongly drive stream insect microdistributions (Mérigoux & Dolédec, 2004). Some aquatic insects choose more exposed surfaces and perform more respiratory movements when flows are low or temperatures high, presumably to increase rates of oxygen supply (Kovalak, 1976; Kovalak, 1979; Wiley & Kohler, 1980; Genkai-Kato et al., 2005). Higher flows can also increase the upper temperature tolerances and lower oxygen limits of stream insects (Frakes et al., 2021).

Here, we examine these ideas in both natural and laboratory settings. We first measured the spatial diversity of temperature, oxygen saturation, and flow velocity within riffles at small spatial scales in several streams in western Montana (USA). We then established experimental gradients of each factor (temperature, oxygen, flow) in the laboratory and measured how strongly nymphs of the giant salmonfly, the stonefly *Pteronarcys californica*, chose among them. Broadly, we expected that when nymphs are subjected to oxygen deficiencies, they should move to ameliorating microclimates (lower temperatures, higher concentrations of oxygen, or higher flows). However, we also predicted that nymphs would make stronger choices among levels of the factors that show strong spatial gradients in nature – thus, we predicted that individual will respond more readily to gradients in flow than to gradients in temperature or oxygen.

## Methods

### Field microclimate measurements

In July, 2019, we sampled water temperatures, oxygen saturation, and flow velocities on the Blackfoot River and Rock Creek in western Montana, USA. The Blackfoot River and Rock Creek have similar hydrology and geomorphology. Both are large, cold-water, cobbled mountain streams fed by snowmelt, precipitation, and groundwater. The Blackfoot River originates in the Flathead Mountains and flows ∼120 km southwest through open sage-prairie, rangeland, and canyons into the Clark Fork River near Missoula, Montana. From its west fork, Rock Creek flows ∼100 km north through the Sapphire Mountains, cutting through rangeland and narrow canyons. It joins the Clark Fork River 40 km upstream of the Blackfoot.

We sampled at 9 sites on the Blackfoot River and 10 on Rock Creek. Sites were separated at relatively even intervals (∼10 km) across the elevation gradients of both rivers (1000-1630 m, and 1090-2060 m, respectively). At each site, microclimates were measured surrounding 15 large cobbles (15-50 cm wide) within a 10 × 5 m quadrant in the middle of riffles. Cobbles were chosen haphazardly every meter along three transects within the quadrant. Transects were separated by 5 m (i.e., one at 0, 5, and 10 m within the quadrant). We measured water temperature and oxygen saturation under the bottom surface toward the rear and under the bottom surface toward the front of each cobble and in the free stream ∼ 3/4 of the way up from the stream bed to the water surface. We measured flow velocity on the top surface, at the leading edge, and tail edge of each cobble as well as in the free stream ∼ 3/4 of the way up from the stream bed to the water surface. Measurements therefore represent different microclimates around and among cobbles. To measure temperature and oxygen saturation, we used an optical temperature and oxygen meter (Pyroscience, OXROB3; Aachen, Germany) connected to a meter (Pyroscience, FSO2-C2). The sensor was fitted into a hollow metal rod, which allowed us to insert the tip into the substrate. At each location, the tip was held in place for 30 s. Flows were measured using a portable velocity meter (Global Water, FP111 Flow Probe; Davison, Michigan, USA).

### Laboratory choice experiments

In the laboratory, we measured the preferences of *Pteronarcys californica* nymphs (salmonflies) for different temperature, oxygen, and flow conditions (see Fig. 1 for schematic of experimental setup and Fig. S1 for photos of a salmonfly nymph and the flow chambers). Salmonflies were collected from the Blackfoot River or Rock Creek with a kick screen (91×91 cm with 1 mm mesh openings) and returned to the University of Montana, where they were transferred to 20-liter buckets filled with de-chlorinated tap water and held in a temperature-controlled incubator (Percival Scientific, I-66LLC8; Perry Iowa, USA). Nymphs were held at ∼ 10 °C and fed cottonwood leaves (*Populus spp.*) from the Blackfoot River or Rock Creek. Leaves were soaked in stream water (i.e., conditioned) for two weeks before being given to the nymphs, which softens them and helps bacterial and fungal biofilms establish on their surfaces, from which salmonflies derive much of their nutrition (Eggert & Wallace, 2007). Water in the buckets was oxygenated and stirred by air directed through air stones. Nymphs were held for a minimum of two days and up to three weeks before experimentation. During this pre-experimental period, there was no mortality and we observed vigorous feeding and frequent molting.

**Fig. 1:**
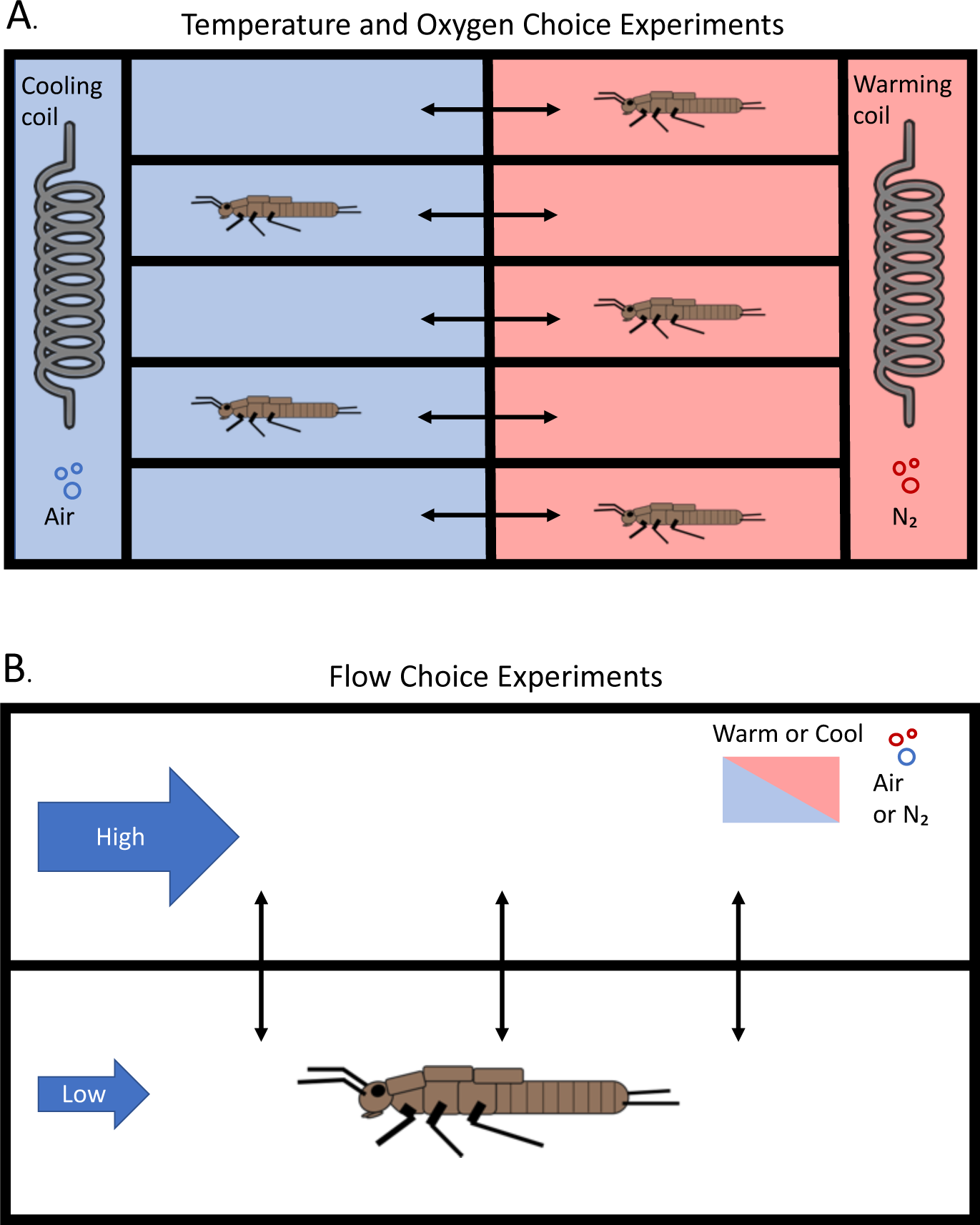
Schematic of temperature and oxygen (A) and flow choice (B) experimental design. In oxygen experiments, nymphs chose between normoxic and hypoxic conditions when temperatures were either cool and warm. In the temperature experiments, nymphs chose between warm and cool conditions when oxygen levels were either normoxic or hypoxic. In the flow choice experiments, nymphs chose between decreasing flows and constantly high flows at variable temperature and oxygen levels.

### Oxygen choice experiments

We performed oxygen choice experiments during the spring and summer of 2019 by measuring where nymphs distributed themselves between two static levels of oxygen saturation (hypoxic versus normoxic) within an experimental aquarium (53×39×16.5 cm). The oxygen gradient was sustained by bubbling air or nitrogen gas through air stones at each end of the aquarium. We placed stainless steel coils connected to cooling recirculating water baths to control the temperature (VWR Scientific, 1160A; Radnor, Pennsylvania, USA & Active Aqua, AACH10HP; Petaluma, CA, USA) and small water pumps to provide flow (PULACO, PL-118, 200 L h^-1^; Guangzhou, China). Mesh walls divided the air stones, coils, and pumps from the compartment in which the nymphs were kept, while still allowing temperature- and oxygen-controlled water to circulate. Plexiglas walls divided the central chamber into five lanes and kept the nymphs separate. Median Plexiglas walls divided the aquarium (and its five lanes) in half, preventing the normoxic and hypoxic water from mixing. However, nymphs were able to move between the hypoxic and normoxic sides by crawling through a 1-cm diameter hole cut in the bottom-center of each median wall. Cobbles and conditioned cottonwood leaves were placed on both sides of all lanes to provide food and substrate for the nymphs. High-grit sandpaper (Gator, P150; Fairborn, Ohio, USA) glued to the floor of the chamber provided footing.

During oxygen experiments, we measured choices made by 30 nymphs – half at 5 °C and half at 16 °C. At the onset of each trial, one nymph was placed on a random side (left or right) of each lane and left to acclimate overnight, during which oxygen levels were held at ∼ 100% of air saturation (∼ 19.0 kPa oxygen at our elevation, 1000 m.a.s.l.) on both sides of the aquarium. On the following morning, the oxygen level on a randomly chosen side of the chamber was decreased to ∼ 30%, 50% or 70% of saturation (∼ 5.7, 9.5, 13.3 kPa, respectively) by bubbling in nitrogen gas, or kept at ∼ 100% of saturation (∼ 19 kPa). Nymphs were left in the chamber for 2-3 hours, after which we recorded their locations (hypoxic vs. normoxic side) and reset the oxygen level on each side of the aquarium to 100% of saturation (19 kPa). Nymphs were then acclimated until the next morning, being free to eat and move between sides of the chamber.

Each nymph experienced every oxygen-choice combination over the course of 5 days. Afterward, each nymph was lighted patted dry with paper towels (Kimtech, Kimwipes; Roswell, Georgia, USA) and weighed. New nymphs were then put in the chamber to experience each choice.

### Temperature choice experiments

During the summer and fall of 2020, we measured how salmonfly nymphs distributed themselves between two static temperatures in an experimental aquarium (49×68×18.5 cm). The arena consisted of two insulated containers (Coleman, 24 Can Party Stackers; Chicago, Illinois, USA) connected via half-inch PVC pipes, providing a route for movement between differing temperatures on either side. Containers were divided into five lanes via mesh walls. Additional mesh walls separated the experimental lanes from the air stones, stainless steel coils, and water pumps that controlled the temperature, oxygen, and flow conditions of each side of the chamber. High-grit sandpaper (Gator, P150) glued to the floor of the chamber provided footing for the nymphs.

We measured temperature preferences of 50 nymphs – 30 in normoxia (100% of air saturation; 19 kPa) and 20 in hypoxia (60 - 95% of air saturation; 11.4 - 18.05 kPa). At the onset of each trial, one nymph was placed on a random side of each lane of the aquarium and left to acclimate for 6 - 12 hours. During this period, oxygen levels were ∼ 100% of saturation (19 kPa) on both sides of the aquarium and temperatures were either 15 °C or 20 °C. For treatments with a 15 °C reference temperature, nymphs were given a choice the following morning between 15 °C versus 15 (control), 20, or 25 °C. These occurred at either normoxia (n = 15) or hypoxia (n = 15). For 20 °C reference trials, nymphs were given a choice of 20 °C versus 20 (control), 15, 25 and 30 °C (normoxia only). Nymphs were left in the chamber for 6-12 hours, after which we recorded the location (warm or cool side) of each nymph and reset the temperature on each side of the aquarium to the reference temperature (15 or 20 °C). Nymphs were then acclimated until the next morning. Each nymph was assigned to either the 15 or the 20 °C reference experiment, but within that experiment they experienced every temperature-choice combination. Once nymphs experienced each choice-trial, we weighed them and placed new nymphs in the chamber for a new round of trials.

### Flow choice experiments

In the flow experiments, nymphs were subjected to ramps of decreasing flow, and we measured the water velocities at which they relocated to higher flows in an adjacent chamber. We conducted these experiments in winter, 2021. The flow chamber (15×17×7.5 cm) was made of glued Plexiglas and divided lengthwise into two lanes by a median wall with a small gap (∼ 0.5 cm high) allowing nymphs to move between sides. High-grit sandpaper (Gator, P150) glued to the floor of the chamber provided footing for the nymphs. Variable flows on the two sides were generated by water pumps (SeaFlo, SFBP1-G500; Xiamen, China) positioned inline in closed loops (PVC) connected to either side of each lane of the chamber via milled nylon end caps (designed in Autodesk based on designs from Mike Nishizaki and milled at The Friday Harbor Laboratories by Adam Summers). Mesh walls inserted between the flow chamber and the end caps prevented nymphs from escaping into the PVC loops. Calibrated flow velocities on each side were controlled by a two-way variable DC power supply (RSR, HY3005F-3; Manheim, PA, USA) connected to the pumps.

We established how flow velocity varied as a function of supply voltage by filming neutrally buoyant microscopic glass particles (TSI, 8-12 μm diameter, 1.5 g/cc; Shoreview, Minnesota, USA) moving through the chamber with a cellphone camera (Samsung Galaxy S10e; Suwon-si, South Korea). A ruler was placed within the chamber in the field of view of the camera during each recording. We used MATLAB (function: DLTdv7) to measure how many pixels spanned 1cm on the ruler, the average pixels the glass beads moved per frame, and the frame rate of the video (MATLAB ver. R2019a). We then calculated the velocity of the water via the following equation: 1/(px/cm) * px/frame * frames/s = cm s^-1^. See Fig. S2 and Table S1 for relationship between power supply voltages, measured water velocies, and interpolated water velocities.

During each experiment, we controlled temperature by submerging the apparatus into a temperature-controlled water bath. Water temperatures were monitored throughout each experiment by a Type-T thermocouple and meter (Barnant, 600-1020; London, Ontario, Canada). Oxygen levels were controlled by bubbling air or mixtures of air and N2 gas into both lanes via small drilled holes on top of the chamber. Oxygen levels were monitored with an oxygen optode (Pyroscience, OXROB3) connected to a meter (Pyroscience FireSting-O2) inserted through an additional hole.

To determine how temperature and oxygen interact with flow to impact microclimate preferences of nymphs, we performed flow choice experiments at two temperatures (10 and 18 °C) and two oxygen conditions (100 and 60% of air saturation; ∼ 19.0 and 11.4 kPa). Ten nymphs were used in each treatment (40 nymphs total). At the onset of the experiment, a nymph was put into a random side of the flow chamber. Preliminary experiments suggested that 9 cm s^-1^ was the maximum velocity in which nymphs could move freely within the chamber without losing their footing and being swept into the mesh at the end. Preliminary experiments also showed that most nymphs explored both sides of the chamber for ∼ 40 minutes before settling into a single lane for at least 2 hours. Based on these observations, nymphs were allowed to acclimate within the chamber for 50 minutes with flow velocities on both sides set to 9 cm s^-1^. Subsequently, we recorded the location (left vs. right side) of the nymph and began to ramp down the flow velocity on that side of the chamber at 10 minute intervals (7, 5, 3, 2, 1.5, 1, 0.75, 0.5, 0.25, and 0 cm s^-1^), while the flow velocity on the other side was kept constant at 9 cm s^-1^. The location of the nymph was recorded at each interval.

### Statistical Analyses

Before analysis, we removed extreme outliers from the microclimate dataset. This was done post hoc by pooling the measurements of each variable and discarding values that were < Q1 – 3 * IQR or > Q3 + 3 * IQR. We used linear mixed-effects models implemented in the R package ‘nlme’ (function: lme) (Pinheiro et al., 2021; R Core Team, 2021) to test whether significant variation in water temperature, oxygen saturation, and flow velocity existed among cobble microclimates (i.e., bottom-fronts, bottom-backs, and tops of cobbles and free stream) and along the length of riffles in the Blackfoot River and Rock Creek. Because we were not interested in conditions at particular sites and cobbles per se, site- and cobble-identify were modeled as random effects. We included the duration of time elapsed from the first sample to each sample per site as a covariate to account for changing conditions throughout the sampling period at each site.

To determine the effect of factor – oxygen, temperature, or flow – on the variability of measurements within riffles and around individual cobbles, we used ANOVA models (function: lm) (R Core Team 2021). We tested the effect of factor on the coefficients of variation (standard deviation / mean) of measurements for each variable at the whole site and individual cobble scales. For each model, river was included as a covariate.

After performing the oxygen choice trials, we discarded data for which oxygen levels on the normoxic side fell below 85% of saturation (16.15 kPa), which was due to equipment malfunction. To analyze effects of temperature and oxygen on salmonfly choices, we used zero-inflated negative binomial mixed models (function: glmm.zinb) in the R package ‘NBZIMM’ (Zhang & Yi 2020). More standard linear mixed effects models could not be used because they resulted in model singularity, likely because the data in many cases were non-normal. NBZIMM uses a non-parametric approach that resolved this issue. Because choices of individual nymphs were measured repeatedly, we used nymph identity as a random effect. In some trials, we were unable to maintain desired temperature and oxygen gradients precisely enough within chambers. To account for this variation, we included the difference between choice conditions (e.g., warm-side temperature – cold-side temperature) on both sides of the chambers as a covariate. In addition, because nymphs in the temperature choice experiment experienced different reference temperatures (i.e., 15 °C and 20 °C), reference temperature was also included as a covariate. For both experiments, preferences for the left versus right side of the chamber in control trials were analyzed by performing exact binomial tests (function: binom.test) (R Core Team).

To test the effects of flow, water temperature, and oxygen saturation on choices nymphs made in flow experiments, we used a mixed-effects logistic regression model, a type of generalized linear mixed-effects model, in the R package ‘lme4’ (function: glmer) (Bates et al., 2007). Because individual nymphs were measured repeatedly, we included nymph ID as a random effect. We also included mass as a covariate. In addition, we analyzed the effects of temperature and oxygen using linear regression models (function: lm) (R Core Team). We first analyzed whether temperature and oxygen levels influenced the water velocity at which nymphs switched between chambers for the final time (i.e., moved to a higher flow). Some nymphs, however, switched to the high flow chamber early during the ramp but later moved back and forth near the end of the trial. This resulted in several low ‘last switch’ flow velocity values, which may have skewed the results of the first model. To account for this potential bias, we also analyzed total time nymphs spent on the high flow side of the chamber.

For all of the above investigations, did not pre-specify a target effect size. None of these experiments has been done previously, and we thus did not know what the variances would be within groups – either for the observational data collected on abiotic conditions within streams or the experimental data on nymph choices made in response to sets of conditions in the lab. Rather, we strived to maximize the number of replicates in each observation or experiment given existing limitations on time and resources.

## Results

### Field microclimate measurements

Oxygen saturation varied little at small scales within riffles of the Blackfoot River and Rock Creek (Fig. 2, Table 1). The mean range within riffles was < 6% of air saturation (1.14 kPa) in both streams. In the Blackfoot, oxygen varied with distance and time elapsed (F_1,111_ = 5.34, P = 0.023 and F_1,111_ = 21.90, P < .001), but not micro-position. In Rock Creek, oxygen varied significantly with micro-position and time elapsed (F_2,272_ = 3.63, P = 0.028 and F_1,137_ = 4.14, P = 0.044, respectively), but not distance. Differences between micro-positions were slight even when significantly different (i.e., < 0.4% of air saturation; 0.08 kPa) and are likely of little relevance to invertebrates, as shown below.

**Fig 2:**
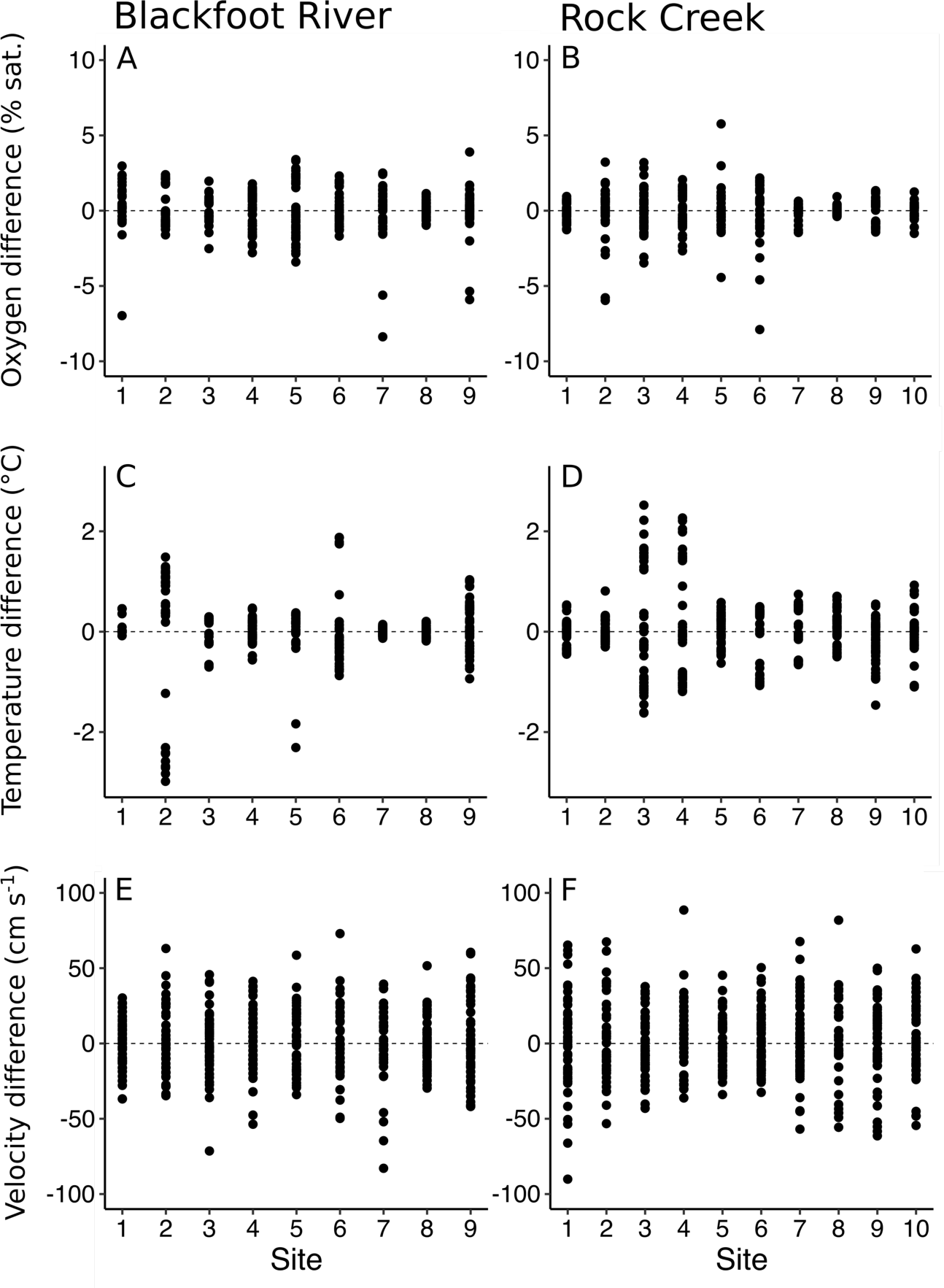
Scatter plots of temperature (A, B), oxygen saturation (C, D), and flow velocity (E, F) samples minus the mean for each site from the Blackfoot River and Rock Creek, respectively. Absolutely values are not reported because they varied strongly with time, and measurements were taken at different days and times.

**Table 1:** Linear mixed effects models for the effects of micro-position, distance along riffles, and time elapsed during sampling at each site on oxygen saturation, water temperature, and flow velocity.

| Oxygen Saturation Microclimates |  |  |  |  |
| --- | --- | --- | --- | --- |
|  | Numerator DF | Denominator DF | F-value | P-value |
| Blackfoot River |  |  |  |  |
| Micro-position | 2 | 231 | 2.23 | 0.110 |
| Distance | 1 | 111 | 5.34 | 0.023 |
| Time elapsed | 1 | 111 | 21.90 | < 0.001 |
| Rock Creek |  |  |  |  |
| Micro-position | 2 | 272 | 3.63 | 0.028 |
| Distance | 1 | 137 | 1.08 | 0.302 |
| Time elapsed | 1 | 137 | 4.14 | 0.044 |

| Water Temperature Microclimates |  |  |  |  |
| --- | --- | --- | --- | --- |
|  | Numerator DF | Denominator DF | F-value | P-value |
| Blackfoot River |  |  |  |  |
| Micro-position | 2 | 235 | 5.07 | 0.007 |
| Distance | 1 | 111 | 36.19 | < 0.001 |
| Time elapsed | 1 | 111 | 1.43 | 0.234 |
| Rock Creek |  |  |  |  |
| Micro-position | 2 | 276 | 6.39 | 0.002 |
| Distance | 1 | 137 | 14.00 | < 0.001 |
| Time elapsed | 1 | 137 | 8.36 | 0.005 |

| Flow Velocity Microclimates |  |  |  |  |
| --- | --- | --- | --- | --- |
|  | Numerator DF | Denominator DF | F-value | P-value |
| Blackfoot River |  |  |  |  |
| Micro-position | 3 | 361 | 232.83 | < 0.001 |
| Distance | 1 | 113 | 0.49 | 0.486 |
| Time elapsed | 1 | 113 | 0.81 | 0.371 |
| Rock Creek |  |  |  |  |
| Micro-position | 3 | 432 | 337.43 | < 0.001 |
| Distance | 1 | 136 | 0.81 | 0.370 |
| Time elapsed | 1 | 136 | 8.49 | 0.004 |

See Table S2 for full tables showing interactions among covariates.

Water temperature varied modestly within stream riffles (Fig. 2, Table 1). The mean temperature range within riffles was ∼ 1.7 °C in both streams. Temperature varied significantly with distance along riffles (Blackfoot: F_1,111_ = 36.19, P < 0.001 and Rock Creek: F_1,137_ = 14.00, P < 0.001), even after accounting for the effect sampling time on micro-thermal variation, which was significant in Rock Creek but not the Blackfoot (F_1,137_ = 8.36, P = 0.005 and F_1,111_ = 1.43, P = 0.234, respectively). On average, temperatures were ∼ 0.7 °C warmer at riffle heads than tails, though they varied more strongly at some sites (e.g., Blackfoot River site 2 where riffle head ∼ 3.2 °C warmer than riffle tail). Micro-position surrounding cobbles also had a significant effect on temperature (F_2,235_ = 5.07, P = 0.007 and F_2,276_ = 6.39, P = 0.002 for Blackfoot and Rock, respectively). However, mean thermal differences between micro-positions were small, particularly between the front and rear of cobbles (i.e., < 0.06 °C), and are likely of little biological importance, as demonstrated below.

Flow velocity varied strongly within stream riffles (Fig. 2, Table 1). The mean range of flow velocities at sites in both rivers were > 125 cm s^-1^. Although flows did not vary along the lengths of riffles (F_1,113_ = 0.49, P = 0.486 and F_1,113_ = 0.81, P = 0.370 for Blackfoot and Rock Creek, respectively), flow velocity depended strongly on micro-position in both rivers (Blackfoot: F_3,361_ = 232.83, P < 0.001; Rock Creek: F_3,432_ = 337.42, P < 0.001). The lowest flows – mean ∼ 20 cm s^-1^ but often near 0 cm s^-1^ – occurred at the front and rear of rocks. Flows were usually high on the tops of cobbles and higher still in the free stream, with a mean of ∼ 64 cm s^-1^ and ∼ 87 cm s^-1^ for both rivers, respectively.

Interactions among covariates were largely non-significant for temperature, oxygen, and flow data in each river and are therefore excluded from Table 1. See Table S2 for all the statistics, including interactions, of each microclimate model.

Factor identity had a significant effect on the coefficients of variation of measurements made within stream riffles (F_2,187_ = 155.03, P < .001) and around individual cobbles (F_2,852_ = 90.26, P < .001) (Table S3). The effect of river was non-significant for each model; interactions were non-significant. Oxygen, temperature, and flow measurements had mean coefficients of variation of 0.01, 0.03, and 0.68, respectively, at the site scale for each river. At the individual cobble scale, oxygen, temperature, and flow measurements had mean coefficients of variation of < 0.01, < 0.01, and 0.32, respectively, for each river.

### Laboratory choice experiments

Oxygen saturation had a significant effect on the microdistributions of salmonfly nymphs in the oxygen choices experiments (F_1,43_ = 16.78, P < 0.001) (Fig. 3A) (Table 2). Choices were strongest when nymphs were exposed to steep oxygen gradients. 100% and ∼ 87% of nymphs were found on the normoxic side when given a choice of 10-35% (1.9 – 6.65 kPa) and 35-60% (6.65 – 11.4 kPa) saturation versus normoxia, respectively. However, when presented with more modest oxygen gradients of 60-85% (11.4 – 16.15 kPa) of saturation versus normoxia, only 45% of nymphs were found on the normoxic side. Initial explorations of the data showed that differences between the oxygen levels on both sides, temperature (Fig. S3A), and nymph mass (range: 0.06 – 1.32g) had no significant effect on oxygen choices, and we the proceeded by pooling the data and dropping these covariates. In addition, there was no significant difference between nymph movements towards the left versus right side of the chamber in the control trials (P = 0.832).

**Fig. 3:**
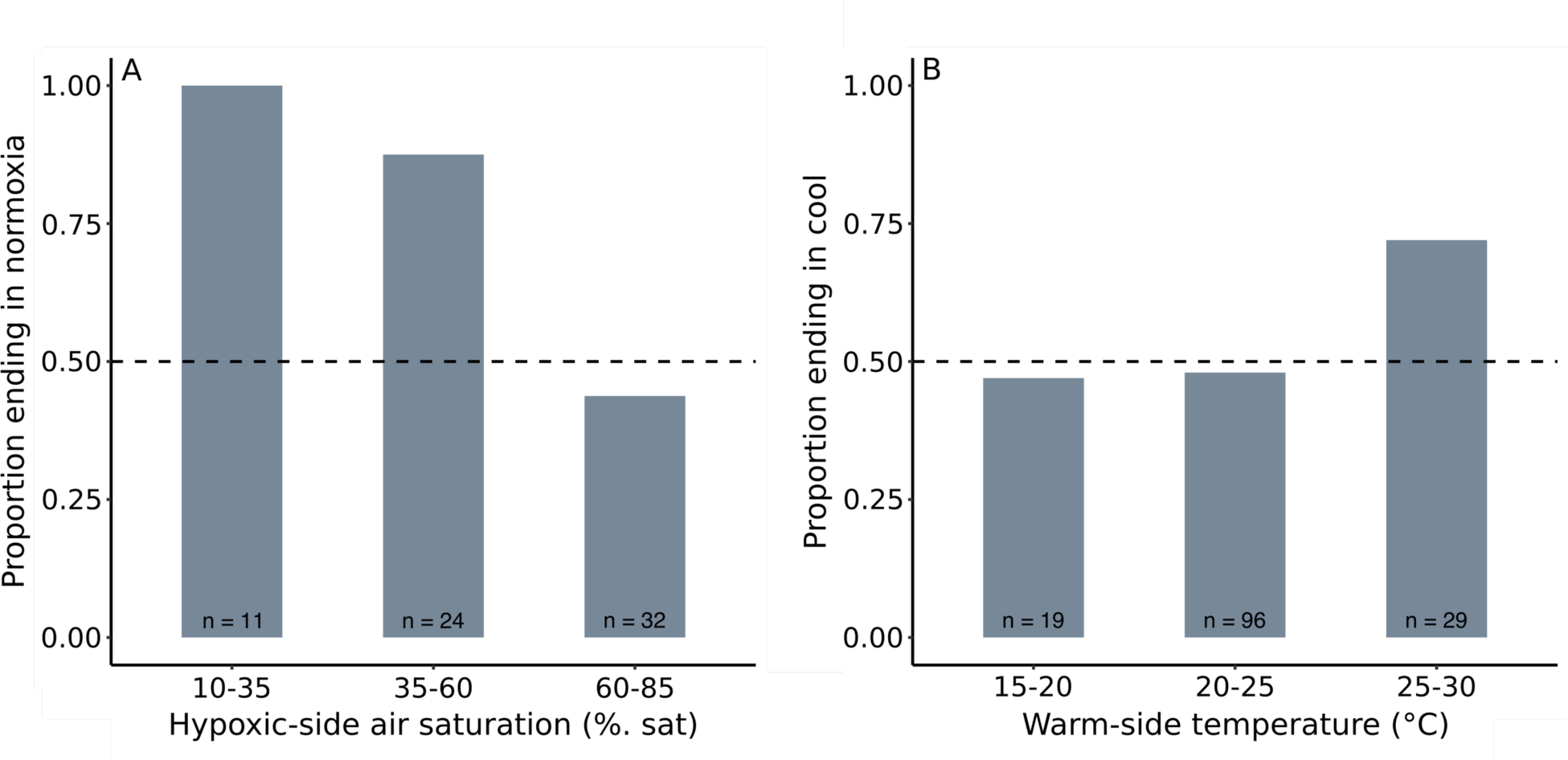
Barplots summarizing choices made by nymphs in oxygen (A) and temperature (B) choice experiments. Nymphs responded strongly to oxygen but, only when presented with strong oxygen gradients. Nymphs made weak temperature-choices and only when presented with extreme thermal gradients, which are likely rare in nature. Because temperature had no effect on oxygen choices and oxygen level had little effect on temperature choices, we do not distinguish the results from the various treatments of each experiment in this figure. See Fig. S6 for more detail. Sample sizes differ strongly across treatments because in some trials, a stronger or weaker temperature or oxygen gradient formed than was intended.

**Table 2:** Zero-inflated negative binomial mixed models for the effect of oxygen and temperature on the location of nymphs in oxygen and temperature choice experiments.

| Oxygen choice experiments |  |  |  |  |
| --- | --- | --- | --- | --- |
|  | Numerator DF | Denominator DF | F-value | P-value |
| Oxygen | 1 | 43 | 16.78 | < 0.001 |

| Temperature choice experiments |  |  |  |  |
| --- | --- | --- | --- | --- |
|  | Numerator DF | Denominator DF | F-value | P-value |
| Temperature | 1 | 77 | 3.55 | 0.063 |
| Oxygen | 1 | 77 | 0.43 | 0.514 |
| Temperature : Oxygen | 1 | 77 | 7.19 | 0.009 |

Temperature had a near-significant effect on distributions of nymphs in the temperature choice experiments (F_1,77_ = 3.55, P = 0.063) (Fig. 3B, Table 2). There was a significant interaction between temperature and oxygen (F_1,77_ = 7.19, P = 0.009), such that nymphs made stronger choices based on temperature when oxygen levels were low (Fig. S3B). However, the effect of this interaction was small, with an interaction coefficient of -0.01. Nymph mass (range: 0.10 – 1.56g), difference between the temperatures on both sides, and reference temperature had no significant effect and were excluded from the final model. There was also no significant difference between nymph movements towards the left versus right side of the chamber in the control trials (P = 0.850).

In the flow choice experiments, flow had a significant effect on the locations of nymphs (z = -4.08, P < 0.001); nearly all nymphs (n = 37/40) moved to the high-flow side before the low-flow side reached 0 cm s^-1^ (Fig. 4, Table 3). The effect of temperature was significant (z = 2.07, P = 0.039), such that nymphs had a stronger preferences for high flows when temperatures were warm. In addition, there was a significant interaction between oxygen and flow (z = -1.99, P = 0.047), such that nymphs relocated to higher flows more readily when oxygen levels were low. Mass (range: 0.18 – 1.40g) had no effect and was dropped from the final model. Supplemental analyses yielded similar results: Oxygen saturation (F_1,36_ = 6.16, P = 0.018) and temperature (F_1,36_ = 5.45, P = 0.025) had a significant effect on the time nymphs spent on the high flow side (Fig. S4A, Table S4); oxygen saturation had a significant effect on the water velocity of the low flow side at the final time nymphs moved from the low to the high flow side of the chamber (F_1,36_ = 11.00, P = 0.002), and temperature had a near significant effect (F_1,36_ = 3.48; P = 0.070) (Fig. S4B, Table S4). No significant interactions were found. Mass (range: 0.18 – 1.40g) had no effect and was dropped from both models.

**Fig. 4:**
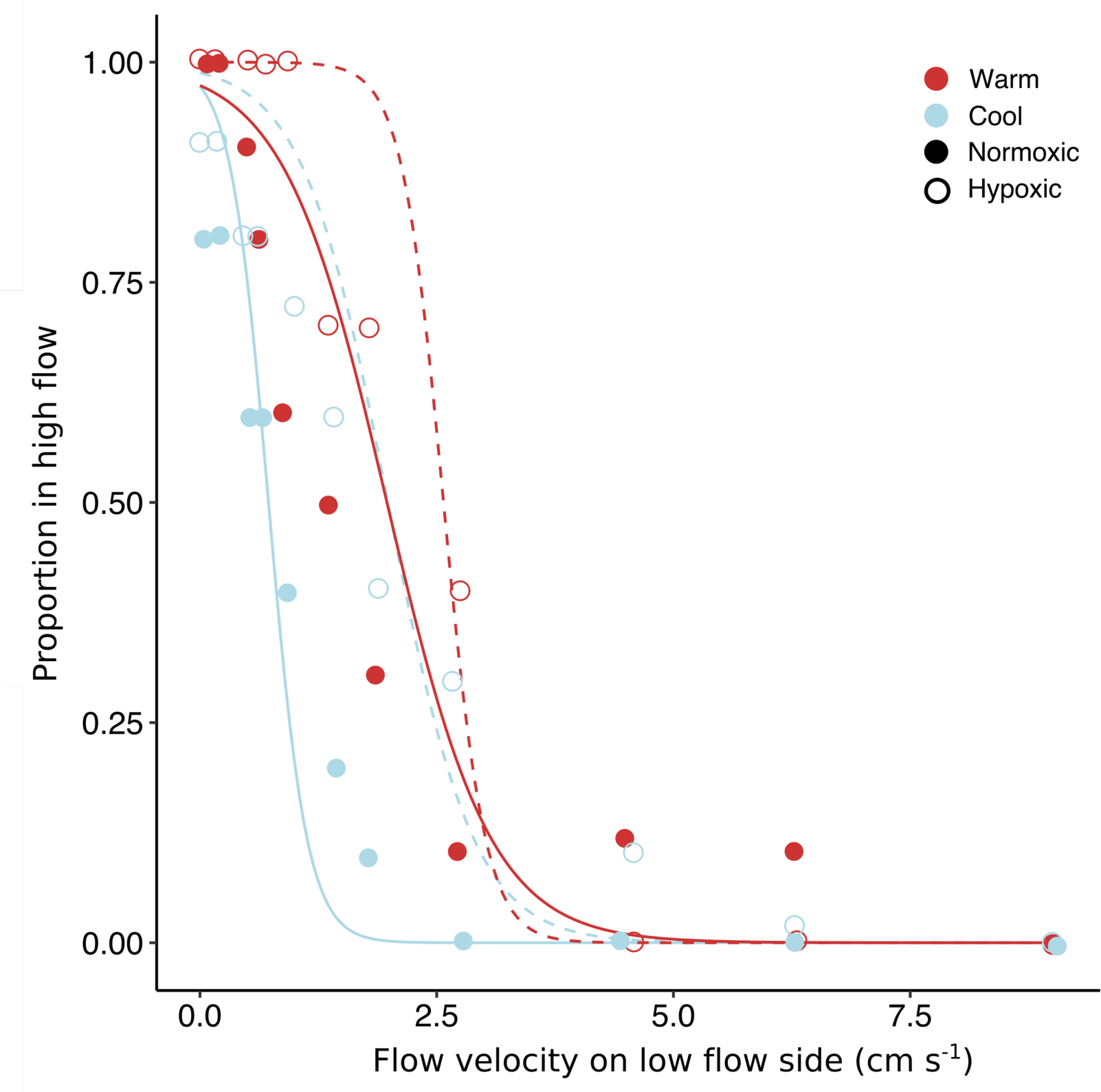
Scatter plot of the proportion of individuals on the high-flow side of the chamber as flows were ramped down on the low flow side during flow choice experiments. The majority of nymphs in each treatment moved to the high-flow side during the ramps. Individuals moved to the high flow side at higher low-flow velocities when exposed to warm or hypoxic conditions. N = 110 for each treatment.

**Table 3:**
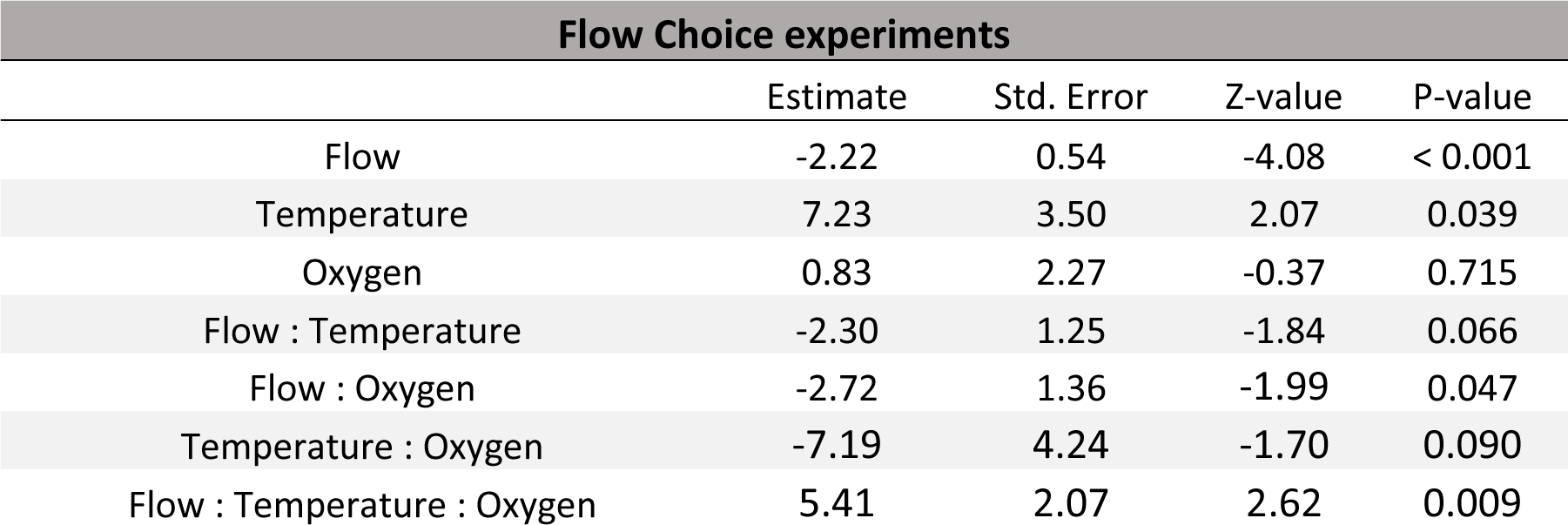
Mixed effects logistic models of the effect of flow velocity, temperature, and oxygen saturation on the locations of nymphs in flow choice experiments

| Flow Choice experiments |  |  |  |  |
| --- | --- | --- | --- | --- |
|  | Estimate | Std. Error | Z-value | P-value |
| Flow | -2.22 | 0.54 | -4.08 | < 0.001 |
| Temperature | 7.23 | 3.50 | 2.07 | 0.039 |
| Oxygen | 0.83 | 2.27 | -0.37 | 0.715 |
| Flow : Temperature | -2.30 | 1.25 | -1.84 | 0.066 |
| Flow : Oxygen | -2.72 | 1.36 | -1.99 | 0.047 |
| Temperature : Oxygen | -7.19 | 4.24 | -1.70 | 0.090 |
| Flow : Temperature : Oxygen | 5.41 | 2.07 | 2.62 | 0.009 |

## Discussion

For insects, aquatic life is challenging because water contains little oxygen (Woods & Moran, 2020; Harrison et al., 2018; Harrison et al., 2012; Verberk et al., 2011). This basic problem of oxygen scarcity can be further exacerbated by warm temperatures and low flows, which increase oxygen demand and decrease oxygen supply, respectively (Verberk et al., 2016; Frakes et al., 2021). For insects, one potential resolution is to exploit mosaics of local microclimates as a way of obtaining combinations of temperature, oxygen, and flow that promote high performance, or that minimize low-oxygen stress (Birrell et al., 2020). Whether insects do so, however, depends on (1) whether biologically meaningful variation in conditions exists at spatial scales accessible by individuals and (2) whether individuals can in fact detect and choose to exploit local variation. Here, we demonstrate that flow velocities in streams varied much more strongly at small scales than did either temperature or oxygen. Accordingly, and in support of our hypothesis, nymphs exploited laboratory gradients of flows far more readily than they did temperature or oxygen; nymphs made choices on temperature and oxygen only when presented with unrealistically strong gradients, which do not reflect field observations.

We measured variation in temperature, oxygen, and flow at two spatial scales in Montana streams – around individual cobbles (scale of ∼ 5 to 50 cm) and along the length of riffles (scale of ∼ 5 to 10 m). Patterns of variation in temperature and oxygen differed at these two scales. At the finest scale – around cobbles, which is most relevant to individual insects – differences were slight: < 0.06 °C and < 0.4% of air saturation (0.8 kPa), respectively. Weak gradients like these offer aquatic insects little opportunity to shift levels of oxygen supply and demand (Verberk et al., 2011) and are probably not actively exploited (as evidenced by the results from our choice experiments). At the larger scale of individual riffles, we found more substantial variation in oxygen and temperature: oxygen saturation varied irregularly by ∼ 6% of air saturation (1.14 kPa), and temperature varied by ∼ 1.7 °C, with water near the tail of the riffle being consistently cooler (∼ 0.7°C) than water near the head. These patterns likely arise because water downwells at riffle heads and is partially replaced by upwelling groundwater at the tails (Davy-Bowker et al., 2006). At several sites, temperatures changed more strongly with distance, being ∼ 3-4 °C cooler at riffle trails. Strong, but inconsistent cold patches have been recorded in other streams in the western US, and are likely important refugia for fish (Ebersole & Frissell, 2001; Ebersole et al., 2003). For stream insects, moving large distances – more than a few meters – likely raises the risks of being dislodged by strong flows and of being predated. Indeed, in a rare mark-recapture study of salmonfly nymphs, Freilich (1991) found that nymphs moved an average of only 1.8 m over a three-month period. In our system, movements at this scale would allow individuals to modify local temperatures and oxygen saturations by an average of ∼ 0.31 °C and 0.11% of saturation (0.02 kPa), respectively. However, drifting can allow stream insects to more easily move greater distances, and may be an important mechanism of exploiting far-away microclimates (Lancaster & Downes 2013).

Flow velocities, by contrast, varied substantially at small spatial scales, often by more than 125 cm s^-1^ around a single cobble. Local flows available to insects are likely even more variable still, as the flow meter we used was too large to fit underneath cobbles, unlike the optode used to measure temperatures and oxygen saturations. We thus could not consistently capture the lowest flows present around cobbles. By moving only a few cm, therefore, individual insects should be able to strongly alter their local flow environments – from < 1 cm s^-1^ under cobbles to > 60 cm s^-1^ on the tops of rocks, decreasing boundary layer thicknesses and increasing rates of oxygen supply (Denny, 1993). Moving greater distances to exploit high flows (e.g., by drifting) is likely ineffective for increasing oxygen supply, as we found that flow velocities did not vary significantly along the lengths of riffles.

In support of our hypothesis that insects should evolve to exploit strong gradients in their local environments, salmonfly nymphs exploited laboratory variation in flow much more readily than they did temperature and oxygen. For example, the temperatures and oxygen levels that induced nymphs to make choices represented gradients that were ∼ 1-2 orders of magnitude greater than those observed in the field. In contrast, nymphs made consistently strong choices in response to ecologically relevant gradients of flow. This suggests that while differences in design between experiments – i.e., two static choices for temperature and oxygen versus a ramp for flow– may have had some influence on the results, nymphs nonetheless exploit local flows more readily than temperature or oxygen. Similar responses to laboratory flows have been observed in mayflies (Wiley & Kohler 1980). In addition, we show some of the first experimental evidence that aquatic insects move to areas of high flow more readily, and spend more time in them, when temperatures are high and oxygen levels low. Our results corroborate field observations of caddisflies and stoneflies, showing that individuals occur more frequently in high-flow microhabitats when bulk flows were low or temperatures high (Kovalak, 1976; Kovalak, 1979; Genkai-Kato et al., 2005). Taken together, these results support the idea that interactions among temperature, oxygen, and flow strongly influence aquatic ectotherm metabolism, performance, and survival (Pörtner, 2007; Rubalcaba et al., 2020; Verberk et al., 2016a b; Harrison et al., 2018; Frakes et al., 2021). In addition, they suggest that aquatic insects exploit mosaics of flow to increase ratios of oxygen supply:demand in nature.

As predicted from the weak gradients in temperature and oxygen measured at the scale of individual cobbles, nymphs made weak choices based on temperature and only when presented with ecologically unrealistic gradients (i.e., 20 vs. 28°C). Nymphs made stronger choices based on oxygen, but as in the temperature experiments, only when gradients were stronger than those found commonly in nature (i.e., 10-55% vs. 100% of air saturation; 1.9-10.5 kPa vs. 19.0 kPa). Counter to our predictions, oxygen choices were not affected by temperature. It’s possible that this is because 15 °C was not warm enough to sufficiently raise metabolic demand of nymphs and elicit a stronger response to oxygen. In addition, although hypoxia led nymphs to make stronger temperature choices, the effect of this interaction was weak (interaction coefficient - 0.01). This is perhaps due to the apparently weak abilities of nymphs to sense and respond to temperature and oxygen. However, they clearly do have at least *some* ability exploit these conditions, which may allow them to avoid stressful conditions that occur irregularly in nature – likely avoiding deep, hypoxic pools or inflows from hot-springs. In more usual circumstances, salmonflies, and other aquatic insects, may rely, at least in part, on their strong acclimation abilities to buffer against warming and hypoxia (Gunderson & Stillman, 2015).

### Conclusions

In the face of climate change and other anthropogenic disturbances, organisms may: go extinct, shift their ranges, evolve new capacities or tolerances, or employ different forms of plasticity (e.g., Doneleson et al., 2019). Our study shows that behavioral plasticity – i.e., moving among locally available flows – may allow stream insects to mitigate oxygen limitation arising from climate change, river dewatering and damming, and nutrient-driven eutrophication (Birrell et al., 2020). Aquatic insects may also rely on other behaviors, like more frequent ventilatory movements, to increase ratios of oxygen supply:demand (Genkai-Kato et al., 2000; Frakes et al., 2021). Other forms of plasticity – i.e., morphological and physiology plasticity – will likely also be important (Angilletta, 2009). For example, aquatic insects like stoneflies and caddisflies tend to grow larger gills (Witchard, 1974; Badcock et al., 1987) and tolerate high temperatures better following acclimation to high temperatures or hypoxia (Gundersen & Stillman, 2015; Malison et al., in review). Indeed, aquatic taxa have about twice the acclimation abilities of terrestrial taxa, which may help buffer them from future functional hypoxia from droughts, warming, and other disturbances (Gunderson & Stillman, 2015).

## Acknowledgements

We thank Michael Nishizaki and Adam Summers for help on constructing the flow choice chamber and Alisha Shah and Alex Birrell for helping measure microclimates in the field. We thank Tim Twardoski, Sean Kellogg, and Elani Borhegyi for assistance in collecting choice data. We acknowledge James Frakes and Jack Hanson for helpful edits on the manuscript. We also thank the Montana Water Center for funding this research through the 2019 Montana Water Center Student Fellowship.

## Data availability

Data will be available from Dryad upon publication.

